# Regulation of Vacuole Fusion in Stomata by Dephosphorylation of the HOPS subunit VPS39

**DOI:** 10.1101/2025.10.02.680005

**Authors:** Anne-Marie Pullen, Grant Billings, Charles Hodgens, Gisele White, Belinda S. Akpa, Marcela Rojas-Pierce

## Abstract

Understanding how plants regulate water loss is important for improving crop productivity. Tight control of stomatal opening and closing is essential for the uptake of CO_2_ while mitigating water vapor loss. The opening of stomata is regulated in part by homotypic vacuole fusion, which is mediated by conserved <u>ho</u>motypic vacuole <u>p</u>rotein <u>s</u>orting (HOPS) and vacuolar SNARE (soluble N-ethylmaleimide sensitive factor attachment protein receptors) complexes. HOPS tethers apposing vacuole membranes and promotes the formation of *trans*-SNARE complexes to mediate fusion. In yeast, HOPS dissociates from the assembled SNARE complex to complete vacuole fusion, but little is known about this process in plants. HOPS-specific subunits VACUOLE PROTEIN SORTING39 (VPS39) and VPS41 are required for homotypic plant vacuole fusion, and a computational model predicted that post-translational modifications of HOPS may be needed for plant stomatal vacuole fusion. Here, we characterized a viable T-DNA insertion allele of *VPS39* which demonstrated a critical role of VPS39 in stomatal vacuole fusion. We found that VPS39 has increased levels of phosphorylation when stomata are closed versus open, and that VPS39 function in stomata and embryonic development requires dynamic changes in phosphorylation. Our data are consistent with VPS39 phosphorylation altering vacuole dynamics in response to environmental cues, similar to well-established phosphorylation cascades that regulate ion transport during stomatal opening.

**SIGNIFICANCE STATEMENT:** Vacuole fusion is important for stomata opening but how it is regulated in response of stomata opening signals is not characterized. This research demonstrated the role of the HOPS complex in vacuole fusion in stomata, and it identified phosphorylation sites in the HOPS subunit VPS39 that are critical for vacuole fusion. One Ser residue was enriched in closed stomata and represents a putative site for control of vacuole fusion downstream of stomata opening signals.

## INTRODUCTION

The Arabidopsis stomatal complex consists of two guard cells surrounding a central pore. Guard cell volume is dynamic and directly influences stomatal pore size, with shrinking leading to closing and swelling leading to opening. The opening and closing of stomata are regulated by movement of water between the apoplast and guard cells. This is largely driven by the movement of ions via transport channels along the plasma membrane and the tonoplast (McAinsh et al. 1990; Andres et al. 2014; Hayashi et al. 2024; Yang et al. 2024). A multitude of signaling pathways must work together to orchestrate stomatal dynamics.

Blue light is a key signal for stomatal opening, and ABA is a key signal for closing. The main pathway controlling blue light-induced stomatal opening involves phosphorylation of PHOTOTROPIN (PHOT) 1 and 2 and the BLUE LIGHT SIGNALING1 (BLUS1) kinase (Takemiya et al. 2013). This ultimately leads to activation of the H^+^ adenosine triphosphatase (H^+^ATPase) pump (Takemiya et al. 2013; Liscum and Briggs 1995; Sullivan et al. 2008), which generates a H^+^ proton gradient and promotes K^+^ and water influx, guard cell swelling, and opening of the stomata (Schroeder et al. 1987). Environmental stressors, such as water deficit, trigger accumulation of Abscisic Acid (ABA), which relays stomatal closing signals through phosphorylation cascades (Schroeder et al. 2001; Kim et al. 2010). In the presence of ABA, PYRABACTIN RESISTANCE/PYRABACTIN-LIKE (PYR/PYL) receptors bind to and inactivate PROTEIN PHOSPHATASE 2C (PP2C) (Park et al. 2009). This allows the OPEN STOMATA1 (OST1) kinase (Geiger et al. 2009; Pei et al. 1997) to activate the SLOW ANION CHANNEL ASSOCIATED 1 (SLAC1) S-type anion channel. Additionally, Ca^2+^ accumulates in the cytosol (Pandey et al., 2007), and anion channels such as SLAC1 HOMOLOGUE 3 (SLAH3) are activated in the presence of Ca^2+^ by calcium-dependent protein kinases (Scherzer et al., 2012). The activity of anion channels SLAC1 and SLAH3 contribute to membrane depolarization. This depolarization activates outward rectifying K^+^ channels like GATED OUTWARDLY RECTIFYING K^+^ CHANNEL (GORK) resulting in K^+^ efflux and water movement out of the cell and into the apoplast (Hosey et al., 2003; Pandey et al., 2007). All of this leads to a drop in guard cell volume and thus stomatal closure (Mustilli et al. 2002). Movement of K^+^ is at least partially controlled by the abundance of K^+^ TRANSPORTER OF ARABIDOPSIS THALIANA 1 (KAT1) at the plasma membrane. The abundance of KAT1 at the plasma membrane is in turn controlled by SYNTAXIN OF PLANTS121 (SYP121) (Leyman et al. 1999; Sutter et al. 2006; Sokolovski et al. 2008; Eisenach et al. 2012). Interestingly, SYP121 function is regulated by phosphorylation in response to light (Ding et al. 2024). Therefore, tight regulation of ion channel activity via phosphorylation is important for multiple stages of stomatal control in response to environmental cues.

Plant vacuoles are important for stomatal function and root cell maturation (Gao et al. 2005; Mirasole et al. 2023; Kruger and Schumacher 2018). Remodeling of vacuole morphology is a main contributor to changing guard cell volume and thus stomatal dynamics (Gao et al. 2005; Cao et al. 2022; Zheng et al. 2014; Mirasole et al. 2023). Guard cell vacuoles are usually fused in open stomata and fragmented and convoluted in closed stomata (Gao et al. 2005; Cao et al. 2022; Zheng et al. 2014). Vacuole fusion is required for full stomatal pore opening (Gao et al. 2005; Zheng et al. 2014). Therefore, understanding the mechanisms controlling vacuole morphology is essential for controlling stomatal movements. In roots, vacuoles exist as highly convoluted networks in the meristematic zone (Scheuring et al. 2024), and homotypic vacuole fusion and fusion of multivesicular bodies (MVB) with the vacuole occur progressively in cells towards the maturation zone (Cajero Sanchez et al. 2018; Viotti et al. 2013; Kruger and Schumacher 2018; Cui et al. 2020). Vacuole fusion is not a passive process but is actively regulated by a suite of vacuole fusion machinery.

Proteins involved in vacuole fusion are highly conserved between plants, animals and yeast (Clary et al. 1990; Wickner and Schekman 2008). In yeast, the homotypic fusion and vacuole protein sorting (HOPS) tethering complex is recruited to the vacuole membrane and holds two membranes in close proximity (Stroupe et al. 2009; Hickey et al. 2009; Brocker et al. 2012; Calakos et al. 1994; Sutton et al. 1998). HOPS acts as an effector for the Rab7 GTPase (Seals et al. 2000; Wang et al. 2003; Wurmser et al. 2000), and Rab7 is needed for HOPS recruitment. HOPS recruits and forms a super complex with vacuolar soluble N-ethylmaleimide sensitive factor attachment protein receptors (SNAREs). At this stage, SNARES are in the trans-SNARE configuration with three Q-SNARES, Qa, Qb, and Qc, at one membrane, and the R-SNARE attached to an apposing membrane (Sutton et al. 1998; Fasshauer et al. 1998). Initial SNARE zippering is followed by HOPS dissociation from the complex and recruitment of Sec17 and Sec18 (Söllner et al. 1993). Sec17 and Sec18 association with the SNARE complex completes fusion and shifts the SNARES into the cis-SNARE conformation (Schwartz et al. 2017; Song et al. 2021).

Plant vacuole fusion is also mediated by conserved HOPS and vacuolar SNARE proteins, but plant-specific functions of these complexes are emerging. The plant HOPS hexamer consists of core proteins VACUOLE PROTEIN SORTING (VPS) 16, 18, 11, 33, and HOPS-specific proteins VPS41, and 39 (Nickerson et al. 2009; Wickner 2010). Unlike in yeast where vacuole traffic and fusion are regulated sequentially via Class C Core Vacuole/Endosome Tethering (CORVET) and later HOPS (Balderhaar and Ungermann 2013), in plants the CORVET complex tethers MVBs to the vacuole while HOPS regulates homotypic vacuole membrane fusion (Takemoto et al. 2018). Moreover, HOPS and CORVET interact with different R-SNAREs in plants with HOPS interacting with VESICLE ASSOCIATED MEMBRANE PROTEIN VAMP713 and CORVET interacting with VAMP727 (Takemoto et al. 2018; Fujiwara et al. 2014; Ebine et al. 2008; Ebine et al. 2011). Plant VPS41 and VPS39 are essential for gametophyte and embryo viability, respectively, and inducible micro-RNA knockdown of either one results in severe vacuole fragmentation in mature root tissue (Takemoto et al. 2018; Brillada et al. 2018). Plant vacuolar SNARE complexes include SYNTAXIN OF PLANTS 22 (SYP22), VACUOLE PROTEIN SORTING 10-INTERACTING 11 (VTI11), SYP51, and VAMP713 or VAMP727 (Ebine et al. 2008; Sanderfoot et al. 2001; Uemura et al. 2004; Takemoto et al. 2018). Mutations in *SYP22* and *VTI11* result in reduced vacuole fusion and slower stomatal opening indicating that SNARE-mediated vacuole fusion is critical for stomatal movements (Zheng et al. 2014; Gao et al. 2005). Gene duplication of SNARE proteins in plants also suggest a diversification of SNARE function (Kanazawa et al. 2016). Overall, conserved and plant-specific regulation of HOPS and SNAREs in vacuole fusion have emerged from plant vacuole fusion studies.

We previously developed a mathematical model to identify key regulatory steps of plant vacuole fusion in stomata based on current understanding from yeast and plants (Hodgens et al. 2024). The model predicted HOPS:SNARE association at the membrane in the stomatal closed state and the existence of a regulatory signal that triggers their dissociation to initiate vacuole fusion. The signal could occur in the form of a post-translation modification (PTM), which could alter the binding affinity between HOPS and SNAREs leading to dissociation of the HOPS:SNARE complex and subsequent vacuole fusion (Hodgens et al. 2024).

Given the prediction of such PTM from the model (Hodgens et al. 2024), the role of phosphorylation in regulating Vps41 and Vps39 function in yeast (Cabrera et al. 2010; Honscher et al. 2014), and the role of phosphorylation in stomatal signal transduction (Ding et al. 2024; Takemiya et al. 2013; Hiyama et al. 2017; Liscum and Briggs 1995; Sullivan et al. 2008; Schroeder et al. 2001; Kim et al. 2010), we hypothesized that a change in phosphorylation state is key to regulation of HOPS function in plant stomata. Here, we first demonstrate the role of the HOPS subunit VPS39 in root and stomatal vacuole fusion using a viable mutant allele of *VPS39*. We present evidence that phosphorylation mutant forms of VPS39 are nonfunctional in both embryo development and stomata, which supports a critical role for VPS39 phosphorylation in vacuole fusion. Moreover, out of all the HOPS and vacuolar SNARE subunits, only VPS39 displays differential phosphorylation between open and closed stomata. These results indicate that cycling between phosphorylation states is essential for VPS39 function and regulation of stomatal vacuole fusion.

## RESULTS

### A viable *vps39* allele displays a mild vacuole phenotype in roots

Knockout alleles of *VPS39, vps39-1* (previously *vps39*) and *vps39-3* (Figure 1A) are not viable which precludes studies of VPS39 function in mature plants (Takemoto et al. 2018). We identified a viable T-DNA insertion allele of *VPS39*, WiscDsLox481-484J4, and named it *vps39-2*. Amplification and sequencing of the flanking region indicated that the T-DNA insertion in *vps39-2* occurred in the 5’ UTR, 2 base pairs upstream of the first codon, and it did not disrupt the *VPS39* protein-coding sequence (Figure. 1A). Transcript quantification by RT-qPCR from whole seedling mRNA showed 2.46 times greater expression of *VPS39* in *vps39-2* compared to the wild type (WT) Col-0 ecotype (Figure 1B), indicating that the T-DNA resulted in altered *VPS39* transcript accumulation. Therefore, the viable mutant is a new tool to study the function of VPS39 in vacuole fusion in vegetative tissues.

**Figure 1.**
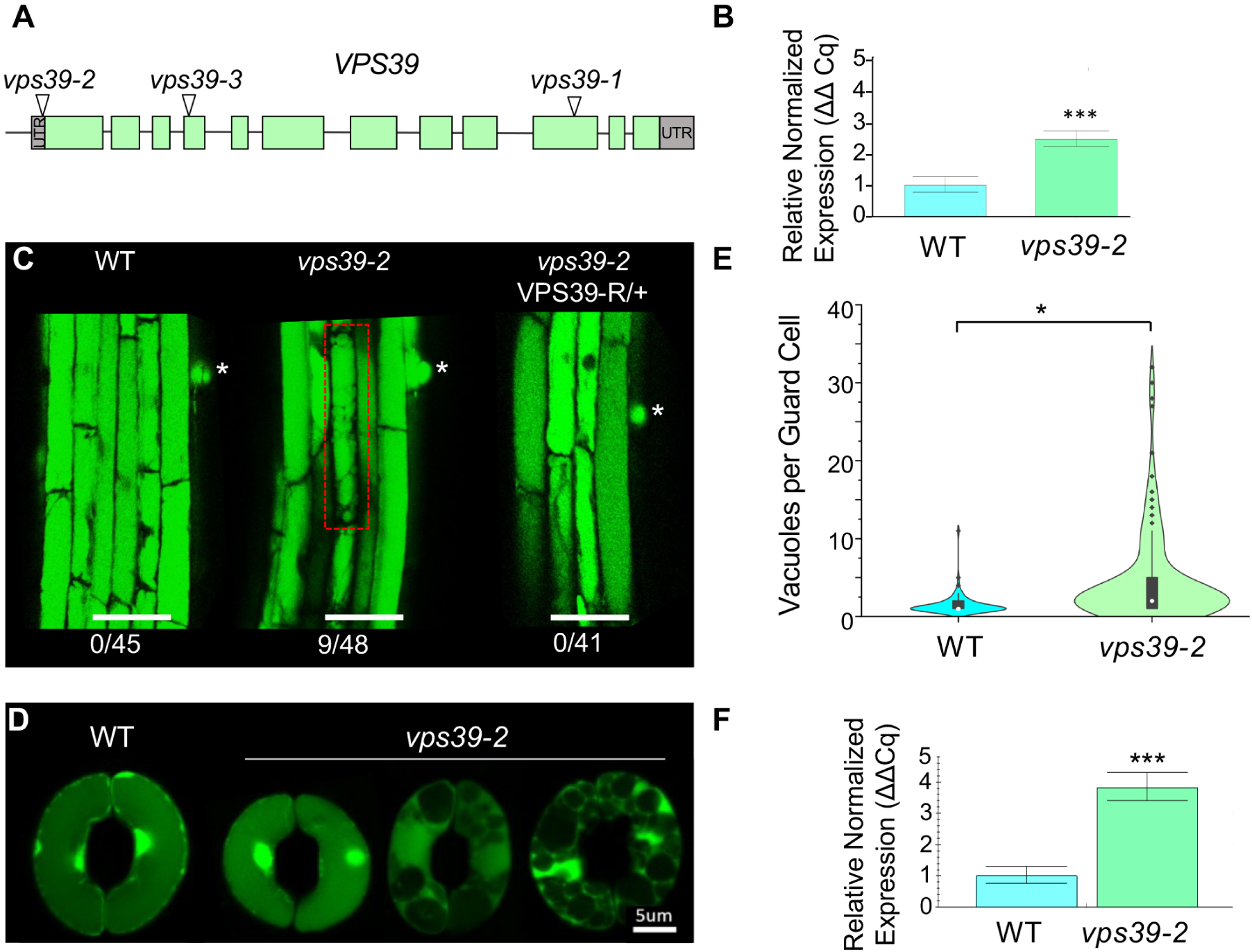
The HOPS subunit VPS39 regulates vacuole fusion in *Arabidopsis*. (A) Map of *VPS39* with the location of T-DNA insertions in *vps39* alleles. Exons represented in blocks and green represents CDS. T-DNA is located 2 bp upstream of translation start site in *vps39-2*. (B) Relative gene expression levels of *VPS39* in 14-day-old seedlings by RT-qPCR. Data presented as relative normalized expression ± 95% confidence interval from n = 30 seedlings per genotype. Gene expression was normalized to *PEX4 (UBC21)* and expressed relative to WT. Significant differences by one-way ANOVA are indicated; p= 6.73x10^-7^ (***). (C) Mild vacuole fragmentation phenotype in *vps39-2* roots. Asterisk represents location of root hair primordia, and a cell with fragmented vacuoles are highlighted in red. The number of seedlings with at least one cell with fragmented vacuoles is shown. Roots were stained with BCECF. Scale bar: 50µm. (D) Vacuole phenotype of *vps39-2* stomata under opening conditions. Leaf epidermal peels of WT and *vps39-2* were exposed to stomatal opening conditions and stained with BCECF. Images of WT and *vps39-2* stomata are shown, with *vps39-2* exhibiting variable vacuole numbers. (E) Vacuole quantification from stomatal opening assays as in (D). A violin plot shows the distribution of the data along with a box and whisker plot with the median noted as a white dot and outliers as black diamonds. Statistical analysis was performed using a generalized linear mixed model comparing vacuoles per guard cell. The model containing genotype was favored (p=0.01). n=97-139 guard cells per genotype from 5-8 plants per genotype. (F) RT-qPCR relative normalized expression of *VPS39* in guard cell-enriched tissue from 5-week-old WT or *vps39-2* plants. n=12 plants per genotype. Data represented as relative normalized expression ± 95% confidence interval. Gene expression was normalized to *PEX4*. Statistical analysis was performed using Mann-Whitney U test; p= 4.66x10^-5^ (***).

To understand how the *vps39-2* mutation affected vacuole dynamics, we first characterized the vacuole maturation phenotype in roots. We took advantage of the morphological gradient of vacuoles from cells in the root meristem to cells in the elongation and maturation zones (Viotti et al. 2013; Kruger and Schumacher 2018). Vacuoles were stained with 2’,7’-Bis-(2-CarboxyEthyl)-5-(and-6)-CarboxyFluorescein, Acetoxymethyl Ester (BCECF) and vacuole morphology was assessed starting from the root transition zone and up to the first root hair bulge. Consistent with previous results, vacuoles in WT seedlings had progressive fusion moving up from the transition and elongation zones with fully fused vacuoles by the first root hair bulge in 100% of seedlings (Figure 1C). In contrast, 18.7% of *vps39-2* seedlings showed a few cells with heavily fragmented rounded vacuoles in elongated and mature root tissue (Figure 1C). Therefore, this mild fragmentation phenotype was not fully penetrant in *vps39-2* seedlings. A complementation analysis showed that expression of a VPS39-RFP fusion driven by the native *VPS39* promoter in the *vps39-2* background restored the WT phenotype with 100% of seedlings having fully fused vacuoles by the first root hair bulge (Figure 1C). Therefore, the mild vacuole phenotype of *vps39-2* is due to the T-DNA insertion, and the mutation may result in a mild decrease in VPS39 protein levels despite slightly elevated levels of *VPS39* transcript. The mild vacuole phenotype of *vps39-2* in roots is consistent with a role of VPS39 in vacuole fusion.

### *vps39-2* mutants have decreased vacuole fusion during stomatal opening

Vacuole fusion is particularly important for stomatal movement where vacuole fusion and fragmentation are concomitant with changes in cellular volume (Gao et al. 2005; Gao et al. 2009; Cao et al. 2022; Zheng et al. 2014). To understand how the mutation in *vps39-2* affected stomata, we conducted stomatal opening assays with epidermal peels of 4-week-old *Arabidopsis* leaves stained with BCECF. Leaf epidermal peels were first treated under stomatal closing conditions for 2 hours and then incubated under light in stomatal opening buffer for 2 hours. Stomata were imaged by super resolution microscopy using Airyscan. As expected (Gao et al. 2005), epidermal peels from WT plants showed mainly one vacuole per guard cell under stomatal opening conditions (Figure 1D, 1E). In contrast, peels from *vps39-2* mutants showed an array of vacuole fragmentation phenotypes ranging from one vacuole per guard cell, to ∼30 vacuoles per guard cell under the same conditions (Figure 1D, 1E). Given the large distribution of vacuole numbers per cell, we used a generalized linear mixed model comparing factors such as date of experiment, individual plant, and genotype (WT vs *vps39-2*) to determine the variable most likely contributing to the fragmentation phenotype. The model containing genotype was favored (p= 0.01) compared to the nested model (containing all other factors except genotype, Data S1). It was clear that the *vps39-2* mutants have a significantly higher number of vacuoles per guard cell compared to the WT. To understand if this phenotype was due to altered expression of *VPS39*, we conducted a RT-qPCR on epidermal peels of 5-week-old Arabidopsis leaves. Similar to the seedling tissue, this experiment showed a 3.68 greater level of *VPS39* transcript accumulation in *vps39-2* when compared to the WT (Figure 1F). Overall, the vacuole phenotype and altered *VPS39* expression in *vps39-2* demonstrate a role of the HOPS complex subunit VPS39 in regulating stomatal vacuole fusion dynamics.

### Phosphorylation is important for VPS39 function

Vacuole fusion in stomata must be responsive to environmental signals to be concomitant with other cellular changes during stomatal opening. A mathematical model predicted that a biochemical trigger that changes the HOPS:SNARE interactions was the main regulatory step for stomatal vacuole fusion (Hodgens et al. 2024). We hypothesized that phosphorylation of a HOPS-specific subunit was important for actuating vacuole fusion in stomata. The Plant PTM Viewer 2.0 database (Willems et al. 2019; Willems et al. 2024) was used to identify possible phosphorylation sites on the HOPS-specific proteins. While 10 phosphorylation sites were reported in VPS39 with high confidence (Mergner et al. 2020; Roitinger et al. 2015; Sugiyama et al. 2008; Umezawa et al. 2013; Wang et al. 2018), only one phosphorylation site, S860, was reported with high confidence in VPS41 (Roitinger et al. 2015; Mergner et al. 2020). We chose three sites in VPS39 for further study, S406, S413 and S806 (Figure 2A, B), because they were detected with high confidence in at least 2 independent studies (Mergner et al. 2020; Roitinger et al. 2015; Sugiyama et al. 2008; Umezawa et al. 2013; Wang et al. 2018). The AlphaFold predicted model of Arabidopsis VPS39 (AF-Q8L5Y0-F1-v4, Jumper et al. 2021; Varadi et al. 2024) shows a β-propeller and an α-solenoid structure typically found in VPS39 and most HOPS subunits from other organisms (Shvarev et al. 2022) (Figure 2A). Two unstructured loops with very low confidence span between AA371-432 and 808-866. VPS39 is annotated by Uniprot (The UniProt Consortium 2025) as having a Citron Homology (CNH) domain (AA16-282), VPS39/Transformation Growth Factor Beta Receptor (TGFβR) association domain 1 (AA502-612), a VPS39-associated zinc finger (AA947-985), and two disordered regions (AA394-414 and 844-864). Intrinsically disordered regions are predicted by <u>MobiDB-lite</u> and Alphafold 2.0 (AA380-429 and 803-867) (Figure 2B) (Necci et al. 2017; Piovesan et al. 2018; Piovesan et al. 2022; Piovesan et al. 2025). Interestingly, S406, S413 and S806 are within or adjacent to the unstructured regions and outside conserved domains. In order to test the importance of these phosphorylation sites for VPS39 function, we generated mutant forms of VPS39-GFP where the three selected residues, S406, S413 and S806, were mutated to alanine (*herein* VPS39^SA^-GFP) or aspartic acid (*herein* VPS39^SD^-GFP), and expressed them under the control of the native *VPS39* promoter. These constructs were introduced into WT, *vps39-2* and *vps39-1* to determine their localization and to test whether they complemented mutant phenotypes.

**Figure 2.**
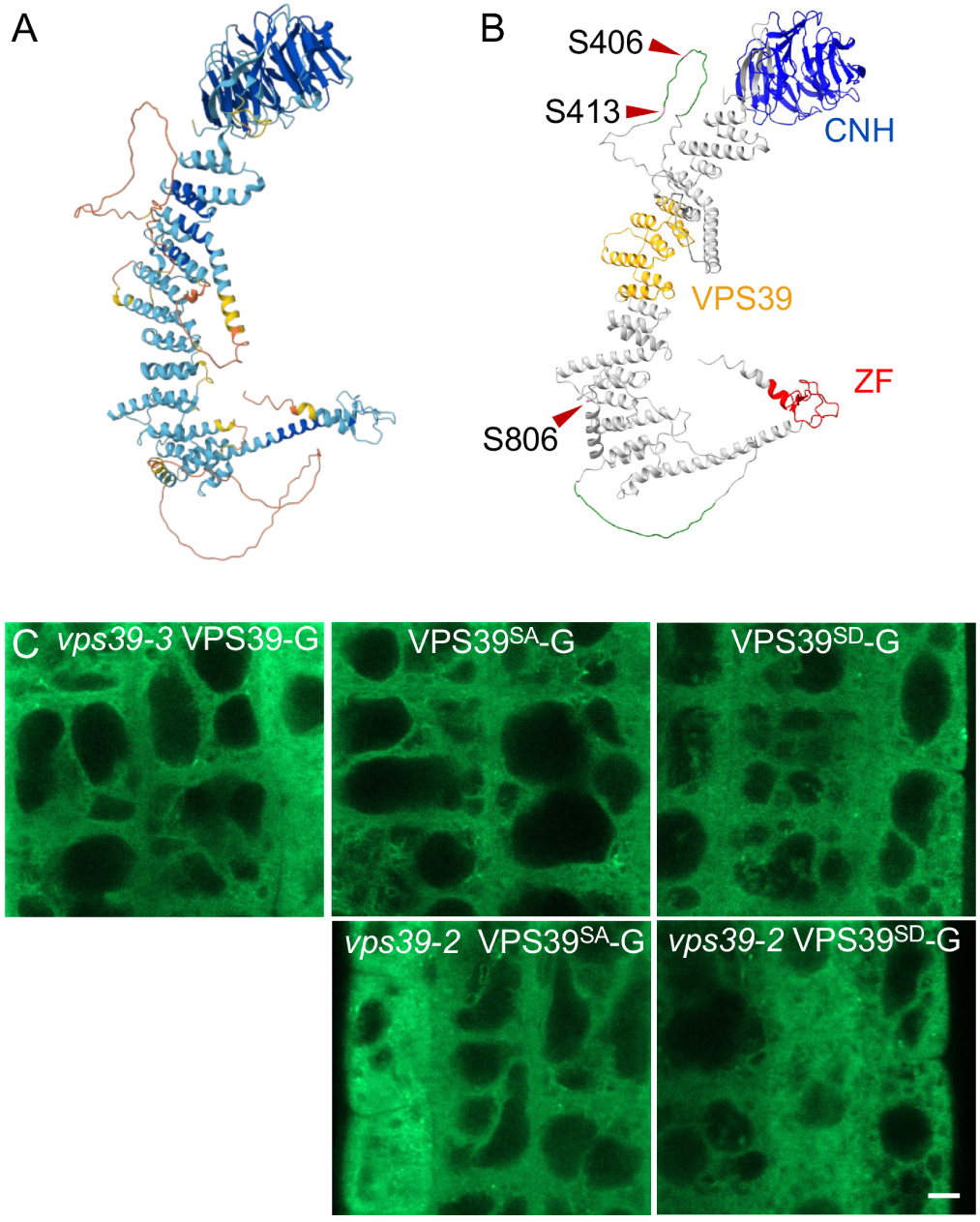
VPS39-GFP phosphorylation mutants do not display altered protein localization. (A) Predicted Arabidopsis VPS39 structure from Alphafold 2.0. (B) Domain structure of VPS39. A Conserved Citron Homology (CNH, blue), a VPS39/TGFβR association domain 1 (orange) and VPS39-associated zinc finger (red) are predicted by Uniprot. Two predicted intrinsically disordered regions (green) are indicated as predicted by MobiDB. The positions of S406, S413, S806 (pink) used for phosphomutant analysis are indicated with arrowheads. (C) Localization of intact or phosphomutant versions of VPS39-GFP in the roots of complemented *vps39-3*, WT, or *vps39-2*. Images are representative from 16-23 seedlings per genotype. Seedlings were stained with BCECF. Scale bar corresponds to 5μm.

We first tested whether the phosphorylation mutants of VPS39 had altered protein localization in roots. Wild type VPS39-GFP localizes primarily to the cytosol, with partial localization to RABG3f-labelled MVBs and minimal enrichment on the tonoplast (Takemoto et al. 2018), which we also observed (Figure 2C). Similar to VPS39-GFP, phosphorylation mutant versions of VPS39 tagged with GFP in the WT background showed largely cytosolic distribution with minimal enrichment on the tonoplast and occasional puncta structures (Figure 2C). Overall, localization of VPS39^SA^-GFP and VPS39^SD^-GFP in the WT and the *vps39-2* backgrounds showed no difference from that of the *vps39-3* VPS39-GFP complemented line (Takemoto et al. 2018) (Figure 2C). These results indicate that phosphorylation does not alter localization of VPS39-GFP in roots at least in plants that express WT copies of *VPS39*.

We then tested whether phosphomutant VPS39 proteins tagged with GFP (VPS39^SA^-GFP or VPS39^SD^-GFP) can complement the embryo lethal phenotype of *vps39-1* knockouts previously described in Takemoto et al. (2018). Homozygous VPS39^SA^-GFP and VPS39^SD^-GFP lines in the WT background were crossed to heterozygous *vps39-1/+* plants, and siliques from F1 plants that were genotyped as *vps39-1/+* VPS39S^A/D^-GFP/+ were imaged under a stereoscope (Figure 3A-C). Embryos from F1 plants should segregate as 1/16 *vps39-1* homozygotes without VPS39S^A/D^-GFP transgene (*vps39-1*/*vps39-1* +/+), 3/16 *vps39-1* homozygotes with at least one copy of the VPS39S^A/D^-GFP transgene (*vps39-1*/*vps39-1* VPS39S^A/D^-GFP/-), and 12/16 having at least one intact *VPS39* endogenous copy regardless of the transgene (*vps39-1*/*VPS39 or VPS39*/*VPS39*). If the phosphomutant VPS39 fusions are functional, only 1/16 aborted embryos should be observed. As previously shown, siliques from heterozygous plants for the *vps39-1* null allele showed segregation of aborted embryos; however, the lethal phenotype of *vps39* nulls is rescued with a VPS39-GFP fusion (Takemoto et al. 2018). The embryo lethal phenotype segregated on average at 2.9% in the WT, 4.2% in the *vps39-3* VPS39-GFP complemented line, 12.7% (∼1/6) in *vps39-1/+* heterozygotes, 10.2 % (∼1/7) in the phospho-null heterozygotes (*vps39-1/+* VPS39^SA^-GFP/+), and 8.75% (∼1/11) in the phospho-mimic heterozygotes (*vps39-1/+* VPS39^SD^-GFP/+) (Figure 3B, Table S1). Therefore, the embryo lethal phenotype did not segregate as 1/16 in either of the phosphomutant siliques, which was confirmed with a chi-square test. A Kruskal-Wallis test followed by a post-hoc Dunn’s test with Šidák correction for multiple comparisons was used to test for significant differences between genotypes. While *vps39-3* VPS39-GFP was not different from the WT, all other genotypes showed significant differences (Figure 3B, Table S1). This result indicates that neither of the phosphomutant versions of VPS39 rescued the loss of *VPS39* function in embryo development. The *vps39-1/+* heterozygotes expressing the phospho-null version of VPS39 (*vps39-1/+* VPS39^SA^-GFP) are not significantly different from the *vps39-1/+* heterozygotes. Nonetheless, siliques from the *vps39-1/+* heterozygotes expressing the phospho-mimic version of VPS39 (*vps39-1/+* VPS39^SD^-GFP) display an embryo abortion phenotype that is significantly different from both *vps39-1/+* and the WT (Figure 3B, Table S1), which suggests a partial rescue. The phenotype of *vps39-1/+* VPS39^SD^-GFP is not significantly different from the complemented mutant (VPS39-GFP *vps39-3*) or from the *vps39-1/+* heterozygotes expressing the phospho-null ( *vps39-1/+* VPS39^SA^-GFP*/+*) (Figure 3B, Table S1). Overall, these results indicate that phosphorylation of VPS39 at either S406, S413 and/or S806 is necessary for embryo development.

**Figure 3.**
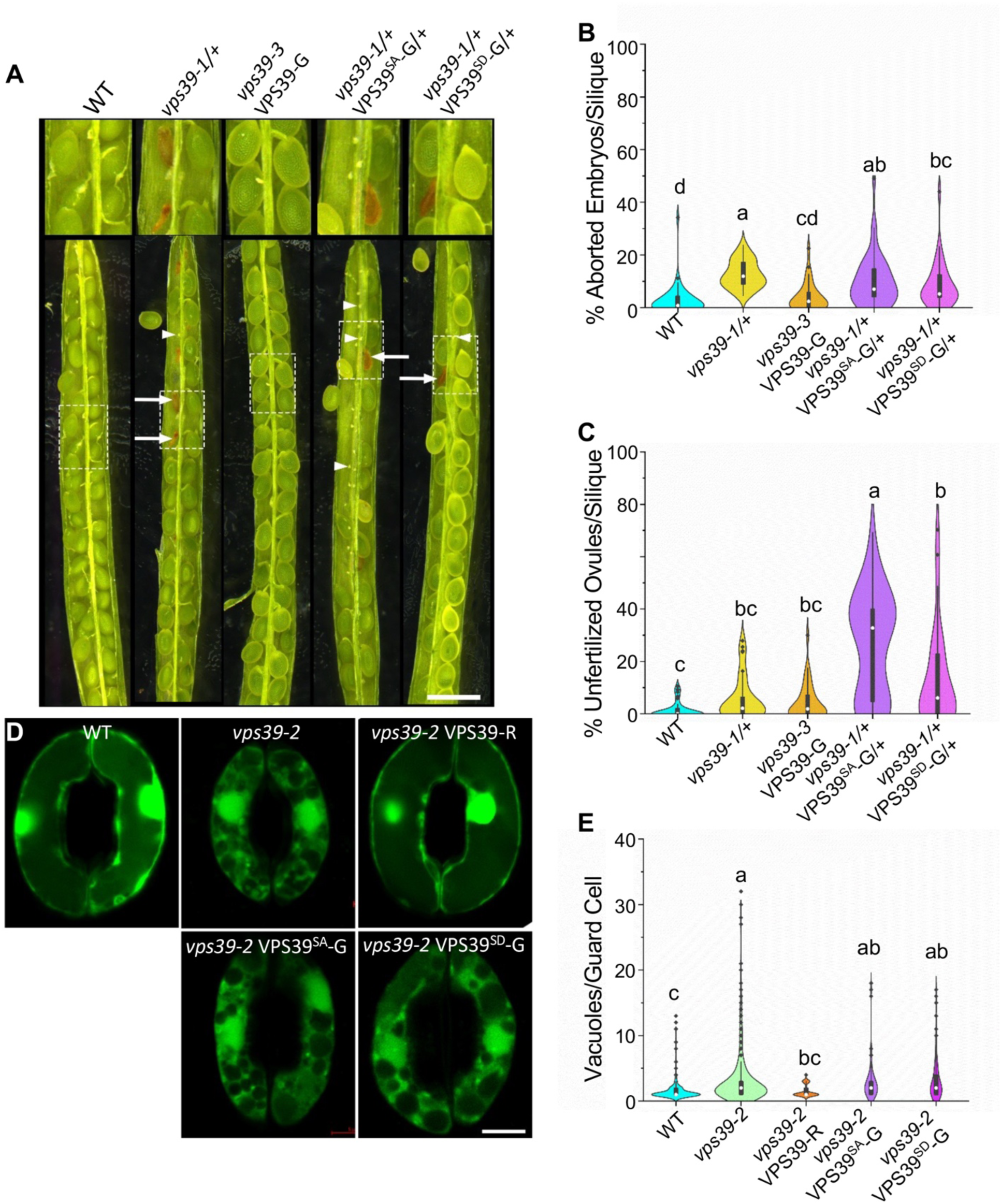
Phosphomutant versions of VPS39-GFP at S406, 413, and 806 are not functional. (A) Embryo abortion phenotype in siliques from WT, complemented *vps39-3* VPS39-GFP, *vps39-1/+* heterozygotes, and *vps39-1/+* carrying the VPS39^SD^-GFP or VPS39^SA^-GFP constructs. Arrows denote aborted embryos. Arrowheads denote unfertilized ovules or very early aborted embryos. Scale bar: 1 mm. (B) Quantification of aborted embryos from (A). Statistical analysis was performed with a Kruskal-Wallis test followed by post-hoc Dunn’s test with Šidák correction. Significant differences shown using compact letter display (p< 0.0051). n= 30-42 siliques/genotype from 6-11 plants/genotype. (C) Quantification of the apparent unfertilized ovules of phosphomutant lines from (A). Kruskal-Wallis followed by post-hoc Dunn’s test with Šidák correction for multiple comparisons p< 0.0051. n= 30-42 siliques per genotype from 6-11 plants per genotype. (D) Vacuole fusion phenotypes of *vps39-2* mutants expressing phosphomutant VPS39-GFP in stomata. Epidermal peels from WT, *vps39-2* and *vps39-2* expressing VPS39^SD^-GFP or VPS39^SA^-GFP were exposed to opening conditions for 2 hours, stained with BCECF, and imaged by confocal microscopy. Examples of highly fragmented vacuoles of phosphomutant lines are shown. Scale bar: 5 µm. (E) Quantification of number of vacuole per guard cell as seen in (D). Statistical analysis was performed using a generalized linear mixed model comparing vacuoles per guard cell followed by bootstrap analysis. Significant difference found in the bootstrap analysis is denoted by compact letter display. The model containing genotype was favored (p= 1.30x10^-7^). n = 97-436 guard cells from 9-26 plants per genotype. This model includes stomata data from Figure 1E. Data distributions in B, C and E are plotted as violin plots along with a box and whisker plots with the median noted as a white dot and outliers as black dots.

Interestingly, we also observed an abundance of unfertilized ovules or early aborted embryos within the siliques of the phosphomutant heterozygotes, which was not observed in the *vps39-1/+* plants (Figure 3C, Table S2). For clarity, these structures will be referred to as “unfertilized ovules”. WT siliques contained an average of 1.5% unfertilized ovules which was not significantly different from the *vps39-1/+* heterozygotes or the complemented *vps39-3* VPS39-GFP line with an average of 5.9% and 4.5%, respectively (Figure 3C and Table S2). In particular, the phospho-null plants (*vps39-1/+* VPS39^SA^-GFP/+) had an average of 26.9% (∼1/4) unfertilized ovules per silique, which was statistically different (α=0.0051) from all other genotypes (Figure 3C, Table S2). The phospho-mimic plants (*vps39-1/+* VPS39^SD^-GFP/+) were unique in that their average and median were quite different. This genotype had an average of 13.3% (∼1/8) unfertilized ovules but a median of 6.0%, which was significantly different from WT but not from the complemented line or *vps39-1/+* (Figure 3C, Table S2). The phosho-mimic line was different from the phospho-null at a p-value of 0.0045 with p<0.0051 denoting significance using the Šidák correction for multiple comparisons (Table S2). These data are consistent with ovule arrest when the *vps39-1* loss of function is combined with the phospho-null *VPS39* transgene (*vps39-1* VPS39^SA^-GFP) in female gametes, a combination that is expected to segregate as ¼ during meiosis. This phenotype could also result from a very early abortion after fertilization. Overall, these results indicate that phosphorylation of VPS39 at S406, S413 and/or S806 is essential for its function during embryo development, and suggests additional roles for VPS39 in the female gametophyte or very early embryo development.

The viable *vps39-2* allele was used to determine the role of phosphorylation on vacuole fusion in stomata. Stomatal opening assays were performed on four-week-old plants with *vps39-2* plants expressing the phosphomutant versions of VPS39 under its native promoter. Levels of vacuole fragmentation under stomatal opening conditions were compared between the WT control, *vps39-2*, a *vps39-2* expressing VPS39-RFP, and the two phosphomutant lines *vps39-2* VPS39^SA^-GFP and *vps39-2* VPS39^SD^-GFP. Consistent with our previous results, WT plants typically showed 1-2 vacuoles per guard cell under these conditions, and the *vps39-2* mutant had a greater number of vacuoles per guard cell compared to WT (Figure 3D, E). The *vps39-2* mutants expressing VPS39-RFP showed in 1-2 vacuoles per guard cell indicating complementation of the vacuole phenotype in stomata. Interestingly, both *vps39-2* lines carrying the phosphomutant VPS39 fusions showed more vacuoles per guard cell compared to WT (Figure 3D, E). A generalized linear mixed model approach was also employed using the variables date, plant and genotype, and the model containing genotype was favored (p= 1.304x10^-7^). Bootstrapped confidence intervals based of the model comparing genotypes (Data S1) determined that there was a difference between the means of the WT and *vps39-2* as well as the WT and *vps39-2* plants expressing VPS39^SA^-GFP or VPS39^SD^-GFP. However, no significant difference was detected between the *vps39-2* background alone and either of the *vps39-2* plants expressing phosphomutant versions of VPS39 ( *vps39-2* VPS39^SD^-GFP or *vps39-2* VPS39^SA^-GFP) (Figure 3E). Consistent with silique data, we do not see a rescue of the *vps39-2* stomata phenotype with the addition of phospho-null or phospho-mimic VPS39. Overall, these data indicate that phosphorylation of VPS39 at S406, S413 and/or S806 is essential for its function during stomatal opening.

### VPS39 undergoes changes in phosphorylation between open and closed stomata

Finding that phosphorylation of VPS39 is essential for its function prompted us to identify changes in phosphorylation in vacuole fusion proteins between open and closed stomata. A global phosphoproteomics study of guard cell-enriched tissue was used to analyze the phosphorylation state of HOPS subunits and vacuolar SNAREs. Epidermal fragments from mature *Arabidopsis* WT leaves were blended and filtered to enrich for guard cells (Jalakas et al. 2017). The fragments containing intact guard cells were then exposed to stomatal opening or closing conditions for 2 hours, and the viability and response of stomata were confirmed in the microscope (Figure S1). Total crude protein extracts were quantified, biological replicates of each respective condition were mixed, and then divided into three standardized technical replicates prior to submission. Samples were subject to phosphoproteomic analysis to determine protein and phosphopeptide levels in each sample. Phosphoproteome values were adjusted using the MSstatsPTM package to normalize each detected phosphopeptide to the amount of the corresponding protein in a given sample (Data S2). Phosphoproteome values discussed below correspond to the adjusted value unless stated otherwise. Differences between open and closed stomata were quantified, and significantly different values were defined as p< 0.05 and Log2FC >0.5 or < −0.5, where a positive Log2FC denotes greater phosphorylation in the closed state and a negative Log2FC denotes greater phosphorylation in the open state. To confirm our treatments and detection methods were accurate, we first verified the expected responses of well-characterized stomatal opening and closing phosphorylation events. Two critical residues of PHOT1, S185 and S350, are phosphorylated in response to blue light (Sullivan et al. 2008), and both showed increased phosphorylation in the open condition with Log2FC of −0.61, and −0.94, respectively. PHOT1 was detected twice with increased phosphorylation in open stomata at S410 at Log2FC of −1.18 and −1.86 (Table S3). Increased phosphorylation of OST1 at S175 (Wang et al. 2023) was also detected in the closed stomata dataset at a Log2FC of 2.21 (Table S3). These results indicate that this is a valid dataset for measurement of protein phosphorylation in open and closed stomata.

The resulting proteome and phosphoproteome datasets were searched for the presence and phosphorylation status of HOPS subunits, vacuolar SNAREs, and CORVET-specific subunits VPS8 and VPS3. Among HOPS and CORVET subunits, all but VPS16 and VPS8 were detected in the proteome (Figure 4A, Table S4). Among these, phosphorylation was only detected in the HOPS-specific subunits, VPS41 and VPS39 (Figure 4, Table S5). The vacuolar SNARE subunits VTI11, SYP22, SYP51, VAMP727, and VAMP711 were also detected in the proteome (Table S4). Among these SNAREs, only VTI11 and VAMP727 were detected as being phosphorylated, which was at S25, and S79 for VTI11 and S194 for VAM727 (Table S5). Neither was differentially phosphorylated between the open and closed samples (Table S5). Therefore, phosphorylation of HOPS-specific subunits appears to be the only regulatory phosphorylation within vacuole fusion machinery in stomata movements.

**Figure 4.**
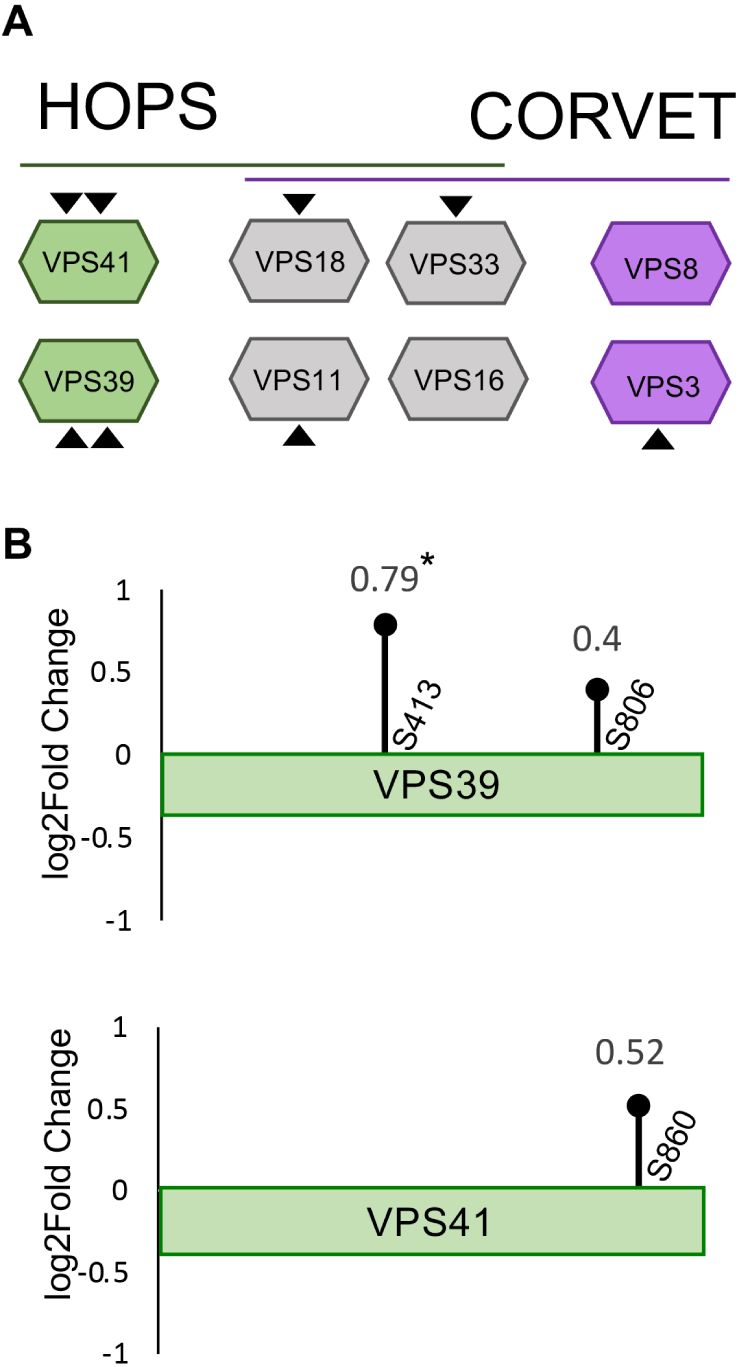
Phosphoproteomics analysis of guard cell-enriched tissue reveals VPS39 phosphorylation changes between open and closed stomata. (A) Depiction of HOPS-specific (green), CORVET-specific (purple) and shared core (gray) subunits detected in the proteome (▴) or the phosphoproteome (▴▴) from guard-cell enriched tissue. (B) Position and Log2FC of phosphorylated residues detected in VPS39 and VPS41 between closed and open samples. Increased phosphorylation was detected in the closed condition for both proteins (Log2FC > 0.5), but only S413 on VPS39 was significant (*, Absolute Log2FC > 0.5 and p<0.05).

Phosphorylation was detected within VPS39 and VPS41. Phosphorylation of VPS39 was detected at S392, S413, and twice at S806 (Figure 4, Table S5), but differential phosphorylation at S413 was the only one that was significant between closed and open states. With a Log2FC of S413 of 0.79 and a p-value of 5.41x10^-6^ (Figure 4B, Table S5), S413 had significantly higher levels of phosphorylation under closed conditions compared to open. Phosphorylation of VPS41 was only detected at S860 with a Log2FC of 0.52 (Figure 4B, Table S5), but was not significant with a p-value of p = 0.08. Therefore, it is not clear whether S860 phosphorylation is involved in opening and closing of stomata (Figure. 4B, Table S5). Overall, phosphopeptide analysis indicates that VPS39 is differentially phosphorylated at S413 between closed and open stomata, and this modification is exclusive to VPS39 among all other HOPS, SNARE or CORVET subunits.

## DISCUSSION

### HOPS complex subunit VPS39 is involved in vacuole fusion in vegetative tissue

The essential nature of VPS39, VPS41 and core subunits VPS16/VCL1 and VPS18 makes investigating the role of HOPS in mature plant tissue difficult (Brillada et al. 2018; Takemoto et al. 2018; Rojo et al. 2001). The viable *vps39-2* allele is a valuable resource in studying VPS39 and HOPS function, but the phenotypes were mild and non-penetrant. The T-DNA insertion in *vps39-2* is 2 base pairs upstream of the *VPS39* transcription start site and includes a truncated left border region. The coding sequence of VPS39 remains unaltered, which explains the viability of this mutant. The pDs-Lox T-DNA contains a the 35S promoter about 2 kb upstream of the T-DNA junction (Woody et al. 2007), and together with other sequences at the T-DNA border derived from the Cauliflower mosaic virus DNA, could act as promoters and transcriptional initiation sites. This is consistent with the observed increase in *VPS39* transcript abundance in *vps39-2.* The transcript, however, is likely to contain a chimeric 5’UTR that includes T-DNA flanking sequences, and this could alter its translation efficiency and reduced levels of VPS39 protein in some cells. The complementation of *vps39-2* with the VPS39-RFP construct is consistent with this hypothesis. Regardless of the exact translational regulation of *VPS39* in these mutants, it is apparent that changes in its expression result in disruption of vacuole fusion.

Several mutations in genes involved in vacuole biogenesis result in abnormal vacuole fusion in stomata. Loss of function alleles for the vacuolar SNAREs VTI11 and SYP22 showed fragmented stomatal vacuoles in response to opening cues (Kato et al. 2002; Zheng et al. 2014; Gao et al. 2005). The viable *vps39-2* allele displayed greater levels of vacuole fragmentation in stomata when compared to the WT. The numbers of vacuoles per guard cell for *vps39-2* are highly variable even on the same leaf. Some of the *vps39-2* plants showed vacuole numbers more similar to WT (1-5 vacuoles per guard cell) while others had much higher numbers (10-30 and beyond). The ability of many of the *vps39-2* guard cell vacuoles to complete fusion indicates that some active VPS39 is present in these tissues. There is also the phenomenon of “stomatal patchiness” wherein sub-regions of stomata on a leaf surface act in a semi-autonomous network to other regions of the leaf countering the generally held belief that all stomata on the leaf surface are homogeneous in their responses (Mott 1995; Peak et al. 2023). The variation in *vps39-2* samples may be due to patchiness of vacuole fusion responses or patchiness of *VPS39* transcription or translation in response to environmental cues. Even WT plants show some level of variation along a leaf; however, it is far less than the *vps39-2* mutants. Without normal regulation of gene expression, even mild changes in available *VPS39* transcript or protein can disrupt vacuole fusion.

### HOPS-specific subunit VPS39 displays changes in phosphorylation between open and closed stomata

A phospho-proteomic analysis was used to identify changes in phosphorylation of proteins involved in vacuole fusion in guard cells. This analysis revealed phosphorylation of VPS39 and VPS41 in intact, non-protoplasted, guard cells from epidermal peels. All of the VPS41 and VPS39 residues detected as phosphorylated in our study have been detected in previous phosphoproteome datasets of different Arabidopsis tissues (Mergner et al. 2020; Roitinger et al. 2015; Sugiyama et al. 2008; Umezawa et al. 2013; Wang et al. 2018; Willems et al. 2019; Willems et al. 2024). Phosphorylation of VPS39 S806 was detected in a study on DNA damage repair in seedlings, but it was not changed by irradiation treatment (Roitinger et al. 2015). Another study detected S806 phosphorylation in 6 of the 30 tissue types being screened in Arabidopsis, although the tissue types were not specified (Mergner et al. 2020). Phosphorylation of VPS39 S413 was detected in 5 studies (Roitinger et al. 2015; Sugiyama et al. 2008; Umezawa et al. 2013; Wang et al. 2018; Mergner et al. 2020). However, S413 phosphorylation was not found in all replicates and/or shown to vary between controls and treatments (Roitinger et al. 2015; Sugiyama et al. 2008; Umezawa et al. 2013; Wang et al. 2018). None of these studies have investigated phosphorylation in guard cell protoplasts or guard cell-enriched epidermal tissue. Our results establish a role for S413 phosphorylation in response to conditions that affect stomatal opening and closing. Additionally, despite the full proteome detecting three core proteins, one CORVET-specific subunit, and some vacuolar SNARE proteins, the only significant change in phosphorylation detected was for VPS39. These results support our hypothesis that the phosphorylation state of VPS39, specifically at S413, changes between stomatal opening and closing.

To understand how these phosphorylation sites could affect VPS39 function, we generated triple phospho-mimic and phosphor-null mutants at S406, S413 and S806 based on previously published phospho-proteomic data (Mergner et al. 2020; Roitinger et al. 2015; Sugiyama et al. 2008; Umezawa et al. 2013; Wang et al. 2018; Willems et al. 2019; Willems et al. 2024). Neither of the phosphorylation mutants complemented the *vps39-1* embryo lethal phenotype or the *vps39-2* fragmented vacuole phenotype. The increased number of unfertilized ovules in *vps39*/+ phospho-null siliques is consistent with aberrant ovules or very early embryo arrest due to the presence of the unphosphorylatable VPS39 (VPS39^SA^) in a null background. Additional studies are needed to determine the exact role of phosphorylation of VPS39 in gamete development and fertilization. Neither phosphomutant VPS39 cassette fully complemented the *vps39-2* allele in stomata, which underscores the importance of VPS39 phosphorylation and dephosphorylation for VPS39 function in vacuole fusion.

The silique and stomata phenotypes of both phosphomutant VPS39 lines indicate that a cycling of the VPS39 protein between a phosphorylated and non-phosphorylated state may be important for its function. Other proteins also lose function when their ability to cycle through their phosphorylation state is disrupted. For example, the FER-LIKE IRON DEFICIENCY-INDUCED TRANSCRIPTION FACTOR (FIT) phospho-mimic and phospho-null mutants at Y278 showed similar loss of nuclear localization compared to WT (Gratz et al. 2020). Additionally, phospho-mimic and phospho-null mutants at S744 and S746 of the plasma-membrane associated protein NON-PHOTOTROPIC HYPOCOTYL 3 (NPH3) showed loss of light-induced relocalization (Sullivan et al. 2021). NPH3 is a regulator of phototropic growth directly phosphorylated by PHOT1. In these examples, cycles of phosphorylation and dephosphorylation state are important for protein function (Sullivan et al. 2021; Gratz et al. 2020).

We propose a working hypothesis for the control of stomatal vacuole membrane fusion by phosphorylation of HOPS (Figure 5). Given that levels of VPS39 phosphorylation at S413 were higher in the closed state, we propose that VPS39 is mostly phosphorylated in closed stomata where vacuoles are fragmented and convoluted. Our results suggest that dephosphorylation of VPS39 is important for triggering vacuole fusion in stomata when opening signals are received, which results in accumulation of dephosphorylated VPS39 in open stomata. Following the model prediction (Hodgens et al. 2024), changes in phosphorylation of VPS39 may result in differential binding affinity between HOPS and the SNARE complex, but this needs to be tested experimentally. Future studies are needed to identify the regulatory phosphatases or kinases involved.

**Figure 5.**
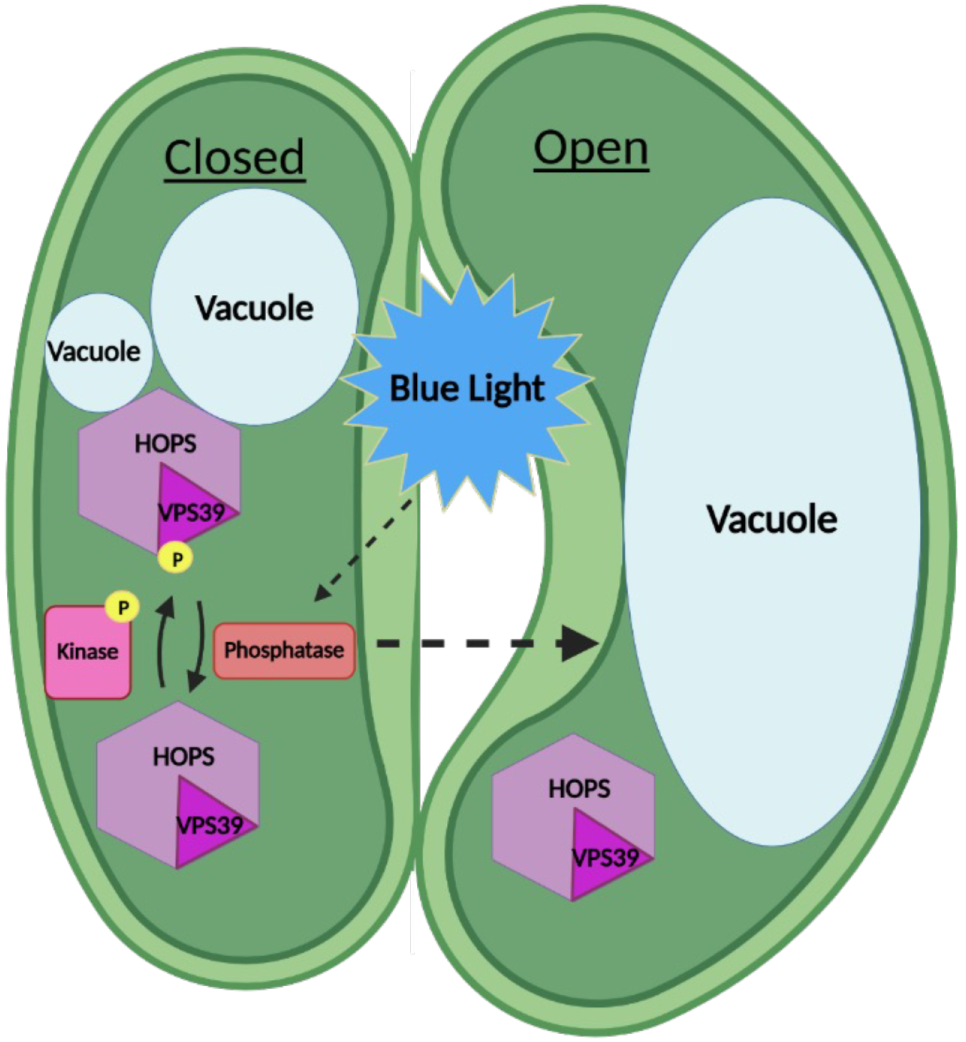
Proposed model for regulation of stomatal vacuole fusion by phosphorylation of HOPS. We predict that the HOPS subunit VPS39 accumulates in its phosphorylated in closed stomata, which may promote HOPS association with the vacuolar SNAREs at the tonoplast. Dissociation of HOPS is necessary to complete fusion, and this could be triggered by VPS39 dephosphorylation in response to stomata opening signals such as blue light. Kinases and phosphatases involved in this process are unknown. Image generated in BioRender (Biorender.com).

### Phosphorylation of VPS39 may regulate vacuole fusion in a plant-specific manner

The morphology of stomata is surprisingly similar throughout plant evolutionary history with only graminaceous monocots deviating from the kidney bean-shaped stomatal complex. Early plant lineages such as mosses and hornwort possess kidney bean-shaped stomata, but these are only found on the sporophyte and are thought to be involved in spore dehiscence (Clark et al. 2022; Chater et al. 2016). The genes for stomata pattering are present in all plant lineages, but it is thought that functional stomata begin to appear in the ferns. Unlike in mosses, some fern stomata can respond to light, ABA, and CO_2_ (Kubasek et al. 2021; Sussmilch et al. 2019; Cai et al. 2017; Cai et al. 2021). It is possible that as land plants evolved and adapted to their new lifestyle, unique ways of regulating vacuole fusion may have arisen in stomata to respond to unique environmental stimuli.

The phosphorylation sites identified in this and previous proteomic studies may point to a plant-specific mechanism of VPS39 phosphorylation in regulating vacuole fusion. These phosphorylation sites lie outside of the conserved domains, and are near intrinsically disorder regions predicted by <u>MobiDB-lite</u> (Piovesan et al. 2025) between amino acid residues 394-413 and 844-864. The relative solvent accessibility score (<u>RSA</u>) of Alpha Fold 2.0 (Varadi et al. 2024) similarly predicts an intrinsically disordered region between residues 380-429, and 803-867 (Jumper et al. 2021; Varadi et al. 2024; Necci et al. 2017; Piovesan et al. 2018; Piovesan et al. 2022; Piovesan et al. 2025). Both predictive models agree that residues S406 and S413 are within intrinsically disordered regions (Figure 2A) (Necci et al. 2017; Piovesan et al. 2018; Piovesan et al. 2022). Additionally, Alphafold2.0 predicted local distance difference test (<u>pLDDT</u>) score measuring the quality of protein structure classified these regions as very low quality (approx. 20-50 out of 100). Accompanying information in AlphaFold presents two reasons for these low scores: (a) the model does not have enough information to fold these regions with confidence. and/or (b) the regions are highly flexible or intrinsically disordered (Jumper et al. 2021; Varadi et al. 2024). Closer evaluation of VPS39 structure from early bryophytes to angiosperms reveal strong conservation within these regions (Figure S2A). VPS39 from *Marchantia polymorpha, Physcomitrium patens, Oryza sativa* and *Arabidopsis thaliana* notably contain the two regions that are predicted to be unfolded/disordered (Figure S2A). While the exact residue may differ between species, it is possible that phosphorylation within these flexible regions contribute to the regulation of VPS39 function across plant lineages. Interestingly the pLDDT protein structure from yeast models display a more tightly folded structure. Though <u>MobiDB-lite</u> (Piovesan et al. 2025) and AlphaFold <u>RSA</u> (Varadi et al. 2024) do predict small regions of disorder within the yeast VPS39 structure, the large unfolded loops found in plant VPS39 are absent in yeast (Figure S2). The differences in predicted structure between yeast and Arabidopsis VPS39, along with the overlap of predicted disordered regions in plants, may point to a plant-specific mechanisms of regulation within these regions.

Changes in phosphorylation regulate HOPS function in yeast; however, very few of the characterized sites for HOPS phosphorylation seem to be conserved between yeast and plants. For example, changes in phosphorylation of the yeast Vps41 amphipathic lipid packing sensor (ALPS) motif is known to regulate its role by switching between tethering of MVBs to the vacuole, or tethering between the vacuole and vesicles derived from the Golgi (Cabrera et al. 2010). However, the ALPS motif, and the kinase that targets it, are missing from plants (Zheng et al. 2014). Alignment of VPS39 predicted structure from Arabidopsis and the one from yeast reveals that the unstructured regions containing S406, S413, and S806 do not occur in the yeast protein (Figure S2B). Moreover, phosphorylation of *Saccharomyces cerevisiae* Vps39 at S246, S247, S249, and S250 regulates its function in vacuole-mitochondria tethering (Honscher et al. 2014), and these phosphorylation sites are located distantly from the Arabidopsis sites in the predicted VPS39 alignment (Figure S2B). Interestingly, the predicted structures for human and Arabidopsis VPS39 align tightly, except for the predicted unstructured loops (Figure S2C). Phosphorylation of human VPS39 has been detected at S441 and T646 (Uniprot), but no function to this phosphorylation has been determined. None of these sites, however, seem to be close to S413 in Arabidopsis. Therefore, phosphorylation at S413 appears to be a plant-specific mechanism for regulation of HOPS function.

Changes in vacuole morphology are key for regulation of stomatal opening (Gao et al. 2005; Zheng et al. 2014; Mirasole et al. 2023), and post-translational modifications of HOPS may be important to control vacuole fusion (Hodgens et al. 2024). We detected higher levels of phosphorylation in VPS39 when stomata are closed versus open, and determined that VPS39 function in embryonic development and stomata requires dynamic changes in phosphorylation. Our data are consistent with VPS39 phosphorylation altering vacuole dynamics in response to environmental cues, similar to well-established phosphorylation cascades that regulate ion transport during stomatal opening. Regions containing phosphorylation sites in the HOPS subunit VPS39 are not conserved between plants and other organisms, and may be important for the stomata to respond to environmental cues. Future identification of kinases and phosphatases involved in VPS39 phosphorylation may identify the precise mechanism of regulation of HOPS by phosphorelay signaling pathways in stomata.

## METHODS

### Plant materials and growth conditions

Arabidopsis ecotype Col-0 was used for all experiments. The *vps39-1* mutant (SALK_092095, *vps39* in Takemoto et al.) and *vps39-2* (CS858015, WiscDsLox481-484J4) (Woody et al. 2007) were acquired from the Arabidopsis Biological Resource Center (ABRC). Complemented *vps39-3* (GABI_376G05) VPS39-GFP and VPS39 RFP lines were previously described (Takemoto et al. 2018). Genotyping of *vps39-2, vps39-1/+* heterozygotes, *vps39-3* and VPS39-GFP fusions was done with primers listed in Table S6. The site of the *vps39-2* T-DNA insertion was determined by Sanger sequencing of flanking fragment amplicons.

Seeds were sterilized and plated on ½ Arabidopsis Growth media (AGM) comprised of 0.2% w/v Murashige and Skoog medium with MES (Research Products International, RPI, M70300-50.0), 1% w/v Sucrose, pH to 5.7 with KOH, and 0.125% w/v GelRite (RPI G35020-250.0). Plates were incubated at 4°C in the dark for 3–5 days before being incubated vertically in a 22°C Percival growth chamber for 5-7 days with ∼120 µmol m^-2^ s^-1^ of light intensity with fluorescent lights (GE 45748) and a 16 h/8 h day/night cycle. Plants used for stomata assays and silique phenotypes were then transplanted to MetroMix 830 (Sun Gro Horticulture) potting mix and grown in a chamber in the NCSU Phytotron under fluorescent lights (Ushio-UFL F32T8/841) providing ∼107 µmol m^-2^ s^-1^ light intensity with 16 h photoperiod, 400 ppm CO_2_ and 22 °C average temperature. Plants used for guard cell (GC) enrichment for phosphoproteomics were grown in potting mix on shelves in the laboratory under LED lights (Monios-L T8 LED Grow Light) at ∼130 µmol m^-2^ s^-1^ of light intensity with 16 h of light. Plants for RT-qPCR of GC-enriched tissue were grown in the Phytotron for 4.5 weeks and moved to the laboratory for 1 week.

### RT-qPCR

Total RNA was extracted from 14-day-old seedlings or 5-week-old guard cell-enriched leaf tissue using RNeasy Plant Mini Kit (Qiagen, 74904) according to manufacturer instructions. This was followed by DNase treatment using DNA Free Turbo DNase kit (Invitrogen AM223). RNA concentration and quality was confirmed using a Nanodrop spectrophotometer for seedling RNA or the Qubit RNA High sensitivity kit (Invitrogen Q32852) for guard cell-enriched samples. Reverse transcription was conducted using GoScript Reverse Transcriptase (Promega A5003). qPCR was conducted using the Bio-Rad CFX Connect real-time PCR system and SYBR Green Master mix (Bio-Rad 1725121). *PEX4/UBC21* (AT5G25760) and *PUX7* (AT1G14570) were used as housekeeping genes for seedling samples, and *PEX4* was used for GC samples (Skiljaica et al. 2022). Three biological replicates and at least three technical replicates were used. A biological replicate consisted of 10 seedlings per genotype for seedling assays, or 8 leaves per plant from 4 plants for guard cell-enriched samples. RT-qPCR primers are listed in Table S6. ANOVA was conducted to determine significance for the seedlings data (n=30 seedlings) with 2.46 times greater expression and 95% CI 2.22-2.73, and a Mann Whitney U test was used for the GC enriched tissue (n=12 plants) with 3.83 times greater expression and 95% CI 3.40-4.29.

### Characterizing *vps39-2* root phenotype

Roots of 3-4-day-old WT, *vps39-2*, and *vps39-2* VPS39-RFP/+ seedlings were stained with 10 µM (2’,7’-Bis-(2-CarboxyEthyl)-5-(and-6)-CarboxyFluorescein, Acetoxymethyl Ester (BCECF, Fisher Scientific B1170) for 2 hours in the dark at room temperature. Roots were imaged using a Zeiss LSM980 confocal microscope using a Plan-Apochromat 20x/0.8M27 objective. BCECF fluorescence was detected using excitation at 488 nm and emission at 499-526 nm. Laser power varied between 0.2-1%. Roots from 3-5-day-old WT (n= 45), *vps39-2* (n= 48), and *vps39-2* VPS39-RFP/+ (n=41) seedlings were analyzed and the experiment was repeated 3 times over a total of 4 days.

### Stomatal opening assays

Leaves from 4-week-old plants were removed in the morning and epidermal peels were immediately generated as described (Behera and Kudla 2013; Li et al. 2013) with modifications. Briefly, a small rosette leaf fragment was applied, abaxial side down, to a coverslip lightly coated with medical adhesive (Skinister Medical Adhesive). All cells but the epidermis were gently scrapped away with a single edge razor. A 1 mm thick silicone isolator (GraceBio #664170) was used to create wells, and peels were immediately stabilized with stomata buffer (10mM MES pH 6.1) (Schroeder et al. 1993). Stomatal closure was induced by incubating peels in closing buffer (10 mM MES pH 6.1, 40 mM malate, 5 mM CaCl_2_) (Schroeder et al. 1993) supplemented with 50 µM ABA (Sigma Aldrich A1049) at 22 °C in the dark for two hours. Next, peels were quickly rinsed once with opening buffer (10 mM MES pH 6.1, 50 mM KCl) (Schroeder et al. 1993), and stomatal opening was induced by incubation with fresh opening buffer supplemented with 10 µM BCECF (Thermo Fisher B1170), 3 µM fusicoccin (Sigma F0537), and 1.6mM Pluronic F-127 (Thermo Fisher P3000MP). Peels were incubated in opening buffer for 2 hours at 22 °C in a Percival growth chamber in the light (∼110 µmol m^-2^ s^-1^). Vacuoles were imaged on a Zeiss LSM880 confocal microscope using a 40x objective (1.2 N.A.) in Airyscan Mode. Guard cells were found first using a 405 nm laser to avoid BCECF bleaching, and BCECF fluorescence-signal was detected using 488 nm excitation and a BP 495-550 emission filter. Laser power was kept at 0.06-0.2%.

For the initial stomata assay comparing WT and *vps39-2* (Fig 1), biological replicates (n) corresponded to 97 guard cells from 5 WT plants and 139 guard cells from 8 *vps39-2* plants (Data S1). The experiment was repeated over 5 days. For the phosphomutant stomata assay, Number of replicates (n) were as follows: 408 guard cells from 23 plants for WT, 436 guard cells from 26 plants for *vps39-2*, 97 guard cells from 9 plants for *vps39-2* VPS39^SD^-GFP, 107 guard cells from 9 plants for *vps39-2* VPS39^SA^-GFP, and 214 guard cells from 9 plants for *vps39-2* expressing VPS39-RFP (Data S1). The Phosphomutant experiment was run on 3 separate days with even numbers of each genotype and a new plant each day. Descendants of 2 independent T1 lines were used for VPS39^SD^-GFP and one T1 line for VPS39^SA^-GFP.

### Molecular cloning and plant transformation

To generate phosphomutant versions of VPS39, 3.8 kb fragments of the *VPS39* genomic sequence spanning −174 bp to +3,696 bp from the ATG start codon, flanked by SwaI and BstBI sites, were de-novo synthesized (Twist Biosciences). Each fragment encodes S to A or S to D substitutions at S406 S413, and S806. Fragments included a C to G point mutation at 3,276 bp from the ATG to overcome a repeating sequence within intron 9. A C to G mutation was introduced at + 2943 in intron 8 to deactivate the BstXI enzyme cleavage site and aid in differentiating mutant clones from WT plasmids. Each fragment was substituted into the full VPS39-mGFP plasmid (Takemoto et al. 2018) using SwaI and BstBI restriction digestion and T4 DNA ligase (Promega M1801). Resulting plasmids correspond to VPS39p:VPS39^SA^-GFP and VPS39p:VPS39^SD^-GFP in the pHGW backbone.

Agrobacterium-mediated plant transformation was achieved by the droplet method modified from Arabidopsis floral dip method (Martinez-Trujillo et al. 2004; Clough and Bent 1998). Agrobacteria was resuspended in transformation buffer (5% sucrose, 0.02% Silwet-77 (Fisher Scientific NC1915706). Siliques and open flowers were removed. A drop of the resuspended culture was placed on flower buds and left in the dark for 24 hours.

### Silique assay

Two independent T1 homozygous lines from phosphomutants (VPS39^SA^-GFP or VPS39^SD^-GFP) were crossed with *vps39-1/+* heterozygotes. Segregating F1 plants were genotyped to identify *vps39-1/+* heterozygotes, and siliques from 5-6-week-old plants were analyzed as described (Feng and Ma 2017). Siliques were imaged on a Leica Thunder for Model Organisms dissecting scope. Data was analyzed using Kruskal-Wallis followed by post-hoc Dunn’s test with Šidák correction for multiple comparisons (p=0.005116) for a two tailed test. Replicates (n) for this experiment are as follows: 42 siliques total from 11 WT plants, 37 siliques total from 9 *vps39-1/+* plants, 31 siliques total from 9 *vps39-3* VPS39-GFP plants, 38 siliques from 9 *vps39-1/+* VPS39^SA^-GFP/+ plants, and 30 siliques from 6 *vps39-1/+* VPS39^SD^-GFP/+ plants.

### Statistical analysis

Normality of data was determined via Shapiro Wilkes test and variance was determined by Leven’s test using the <u>Origin</u> software package (OriginLab Corporation). A Mann-Whiteny U test at alpha =0.05 was used where one or both assumptions were violated and one comparison was needed. A Kruskal-Wallis followed by post-hoc Dunns test and Šidák correction for multiple comparisons was used when multiple comparisons were required and normality and/or variance assumptions were violated. In cases where subtle phenotypes and high variability were observed (e.g. stomatal vacuole phenotype), a Poisson truncated distribution was constructed, and a generalized linear mixed model was used to determine the effect of genotype on the dataset. Pairwise comparisons between genotypes were made using a bootstrap confidence interval based of the generalized linear mixed model. This was performed using R (4.4.1) in R Studio (R Core Team 2021). Specific packages for the guard-cell vacuole analysis included <u>tidyverse</u> (https://www.tidyverse.org/ (Wickham et al. 2019), <u>lme4</u> (Bates et al. 2015), <u>glmmTMB</u> (McGillycuddy et al. 2025; Brooks et al. 2017), <u>emmeans</u> (Searle et al. 1980), and <u>bootstrap</u> (Davison and Hinkley 1997).

### Guard cell enrichment

Guard cell enrichment was conducted as outlined in Jalakas et al. (2017). A total of 144 WT Col-0 Arabidopsis plants were grown in individual pots to 4-weeks-old for phosphoproteomics. Plants were grown over 3 separate planting periods. Rosette leaves were excised, their mid-ribs were removed with scissors and fragments were kept in sterile ice-cold ddH_2_O water until processing. The leaf fragments were then blended (Oster Blender Pro 1200) with 4 °C sterile ddH_2_O water and ice. Samples were strained through 100 µm nylon mesh (Elko Filtering Co 06-210/33) to collect the epidermal fragments containing intact guard cells. For RNA extraction, samples were immediately flash frozen in liquid nitrogen. For phosphoproteomics, the intact guard cells were then equilibrated with stomata buffer in the dark for 30 minutes, rinsed with sterile ddH_2_O water, and then incubated with opening buffer in the light or closing buffer in the dark at 22 °C for 2 hours. To confirm stomatal opening or closing, a small aliquot of each extraction was mounted on slides and imaged using a compound microscope (Leica DM5000B and DMC4500 camera) using bright field with a 20x water objective. Closed aliquotes were kept in the dark and very quickly imaged at low light. Samples were then washed in sterile ddH_2_O water and flash frozen with liquid nitrogen for protein extraction (Jalakas et al. 2017).

### Protein extraction and quantification

Frozen tissue was ground with liquid nitrogen with mortar and pestle. Proteins were solubilized in 8 M urea (Thermo Fisher 29700), 50 mM Tris–HCl pH 7.8 supplemented with cOmplete™ EDTA-free Protease Inhibitor cocktail (Sigma-Aldrich 11836170001) and Pierce™ Phosphatase Inhibitor Mini Tablets (Thermo Fisher A32957). Lysates were kept on ice for 10 min, mixed by gentle vortexing, and clarified by centrifugation at 8,000 × *g*, 4 °C for 10 min. The supernatant was spun again at 12,000 × *g*, 4 °C for 5 min. Clarified extracts for each condition (“open” or “closed”) were pooled to maximize concentration, partitioned into three technical replicates and snap-frozen. Protein concentration was assessed by Bradford assay (Alfa Aesar / VWR AAJ61522-AP).

### Proteomics and phosphoproteomics

Protein analysis was completed by the UNC Michael Hooker Metabolomics and Proteomics Core. Protein samples were acetone-precipitated, and pellets were resuspended in 1 M urea. Samples were then reduced with 5 mM dithiothreitol (Thermo Fisher 20290) for 45 min at 37 °C, and alkylated with 15 mM iodoacetamide (Thermo Fisher A39271) for 30 min in the dark at room temperature. Samples were digested with Lys-C (Wako/VWR 125-02543; 1:50 w/w) for 2 h at 37 °C, followed by overnight trypsin digestion (Promega V5111; 1:50 w/w) at 37 °C. Peptides were acidified, desalted with desalting spin columns (Thermo Fisher 89852), dried by centrifugation, and quantified with a Pierce Quantitative Colorimetric Peptide Assay (Thermo Fisher 23275).

Six samples (3 open, 3 closed) plus four pooled references (equal-volume mixtures of each sample) were labeled with TMT10 reagents (Thermo Fisher 90406) at room temperature. Labeling efficiency was verified by LC-MS/MS prior to quenching; After confirming exceeding 98% incorporation, reactions were quenched with 0.4% (final concentration) of hydroxylamine (Sigma-Aldrich 467804), combined 1:1, desalted with desalting spin columns (Thermo Fisher 89852), and dried by centrifugation. TMT-labeled peptides were fractionated by high-pH reversed-phase HPLC (Agilent 1260) with a Zorbax 300 Extend-C18 column (3.5 µm, 4.6 × 250 mm) over a 90-min gradient using 4.5 mM ammonium formate (pH 10) in 2% (mobile phase A) or 90% (mobile phase B) (v/v) LC-MS-grade acetonitrile (Sigma-Aldrich 271004-100), essentially as described (Klomp et al. 2024; Mertins et al. 2018). Ninety-six primary fractions were collected and concatenated non-contiguously into 24 fractions; 5% of each was dried and stored at −80 °C for proteome-level analysis. The remaining material was concatenated further into three fractions, dried by vacuum centrifugation and subjected to Fe-NTA enrichment using the High Select Fe-NTA kit (Thermo Fisher A32992) according to the manufacturer’s instructions. Eluates were then dried and stored at −80 °C. For total proteome injections, 205 µg tryptic peptides were allocated per sample and dried down.

Proteome (24 fractions) and phosphoproteome (3 Fe-NTA fractions, analyzed in duplicate) samples were analyzed on an Ultimate 3000 system coupled to an Orbitrap Exploris 480 operating in TurboTMT mode (Thermo Fisher). Peptides were separated on an Ion Opticks Aurora C18 column (75 µm ID × 15 cm, 1.6 µm) at 250 nL/min using 0.1% formic acid in water (mobile phase A) and 0.1% formic acid in 80% acetonitrile (mobile phase B). Gradients were 70 min (proteome) or 100 min (phosphoproteome) from 5–42% B. MS1 scans (m/z 375–1400) were acquired at 60,000 resolution with standard automatic Gain Control (AGC) and maximum injection time set to auto. Data-dependent MS2 scans were acquired at 30,000 resolution (HCD 38%; isolation window 0.7 Da; first mass 110 m/z; AGC target 300%; injection time auto) with a 3-s cycle time.

### Database search and statistical processing

Raw data was processed using Proteome Discoverer (V3.1, Thermo Fisher) set to ‘reporter ion MS2’ with ‘10pex TMT’. Spectra were searched using Sequest HT against the *Arabidopsis thaliana* UniProt reviewed proteome (∼16,000 sequences) appended with common contaminants. Enzyme specificity was trypsin with up to two missed cleavages. Static modifications: carbamidomethyl-Cys and TMT10 on peptide N-termini and Lys. Variable modifications: Met oxidation and protein N-terminal acetylation; phosphorylation on Ser/Thr/Tyr was included for phosphopeptide searches. Precursor and fragment tolerances were 10 ppm and 0.02 Da, respectively. Peptide-spectrum matches were filtered to 1% FDR. Quantification used MS2 reporter ion intensities with a ≤30% co-isolation interference threshold.

Downstream analysis was performed in R 4.4.1 using the MSstatsTMT and MSstatsPTM package (Kohler et al. 2023)(Huang et al. 2020). Spectrum-level intensities underwent global median normalization and were summarized to proteins by Tukey’s median polish. Imputation required ≥1 non-missing TMT channel for the same PSM within a run and ≥1 non-missing PSM from the same protein in the same channel/run. Linear mixed-effects models were used for inference with Benjamini–Hochberg adjustment of p-values. For phosphosite-level results, abundances were adjusted for protein-level changes using MSstatsPTM.

#### Accession numbers

Sequence data from this article can be found in the GenBank/EMBL data libraries under accession number AT4G36630 for Arabidopsis VPS39. Uniprot protein accessions used for protein folding analysis in AlphaFold of VPS39 include Q8L5Y0 for Arabidopsis, A0A2R6W3Y9 for *M. polymorpha*, B8AQN8 for *Oryza sativa*, A0A2K1JXU4 for *P. patens*, Q07468 for yeast and Q96JC1 for human.

## DATA STATEMENT

The data that supports the findings of this study are available in the supplementary material of this article.

## ACKNOWLEDGEMENTS

This work was supported by the National Science Foundation (MCB-1918746 to M.R.P. and B.S.A.), by the Research Capacity Fund (HATCH), project award no. 7005574 (to M.R.P) from the U.S. Department of Agriculture’s National Institute of Food and Agriculture, and in part by NCSU through access to the Cell and Molecular Imaging Facility. We thank the Michael Hooker Metabolomics and Proteomics (MAP) Core at UNC Chapel Hill for proteomics services. We thank Takashi Ueda from the National Institute for Basic biology (Okazaki, Japan) for sharing materials and other members of the Rojas-Pierce lab for helpful discussions.

## SUPPORTING INFORMATION

**Figure S1. Stomata from epidermal fragments used for phosphoproteome.** Bright-field image of epidermal fragments after treatment in opening or closing conditions.

**Figure S2. Chimera X protein folding alignments of VPS39.** A) AlphaFold predictions and alignment of VPS39 orthologs from *Marchantia polymorpha, Physcomitrium patens*, *Oryza sativa,* and *Arabidopsis thaliana*. All plant VPS39 structures are predicted to have two large unstructured loops similar to Arabidopsis. (B-C) Neither *Sacharomyces cerevisiae* or human VPS39 proteins show similar unstructured domains as Arabidopsis.

**Data S1.** Confidence intervals for bootstrap analysis from Generalized linear mixed models for stomata assays

**Data S2.** Proteome and phospho-proteome data for proteins discussed in this study.

**Table S1.** Comparison of medians of aborted embryos from Figure 3B.

**Table S2.** Pairwise comparison of medians for unfertilized ovules from Figure 3C.

**Table S3.** Reference phosphopeptides for PHOT1 and OST1.

**Table S4.** HOPS, CORVET and vacuolar SNARE proteins detected in proteome.

**Table S5.** Detected phosphorylation of HOPS, CORVET and vacuolar SNARE proteins.

**Table S6.** Primers used in this study.

