## Supplemental Material for "Regulation of Vacuole Fusion in Stomata by Dephosphorylation of the HOPS subunit VPS39"

### SUPPLEMENTAL INFORMATION

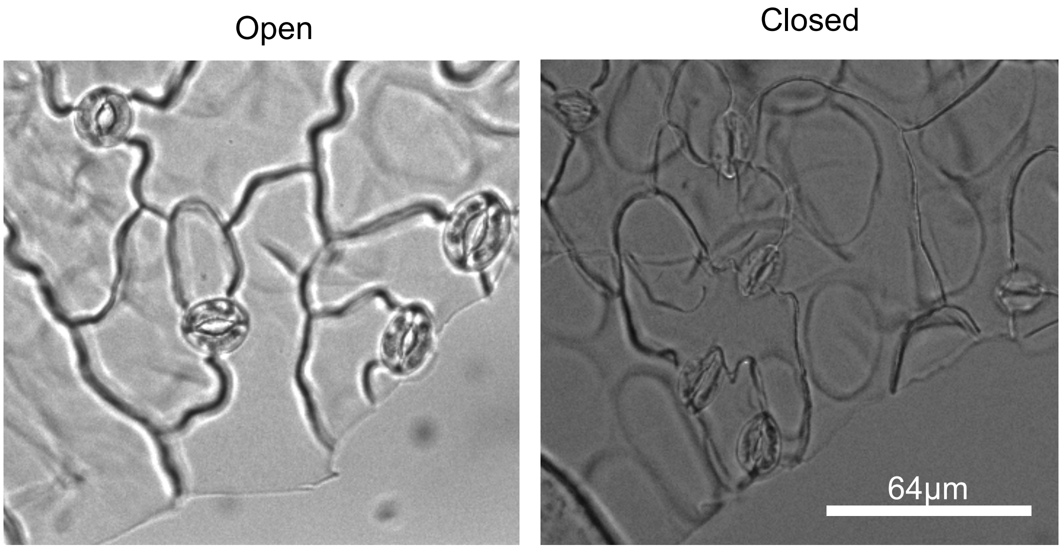


**Supplemental Figure 1. Bright-field image of epidermal fragments used in proteome and phosphoproteome.** Bright-field image of epidermal fragments after treatment in opening or closing conditions as used for phosphoproteomics.

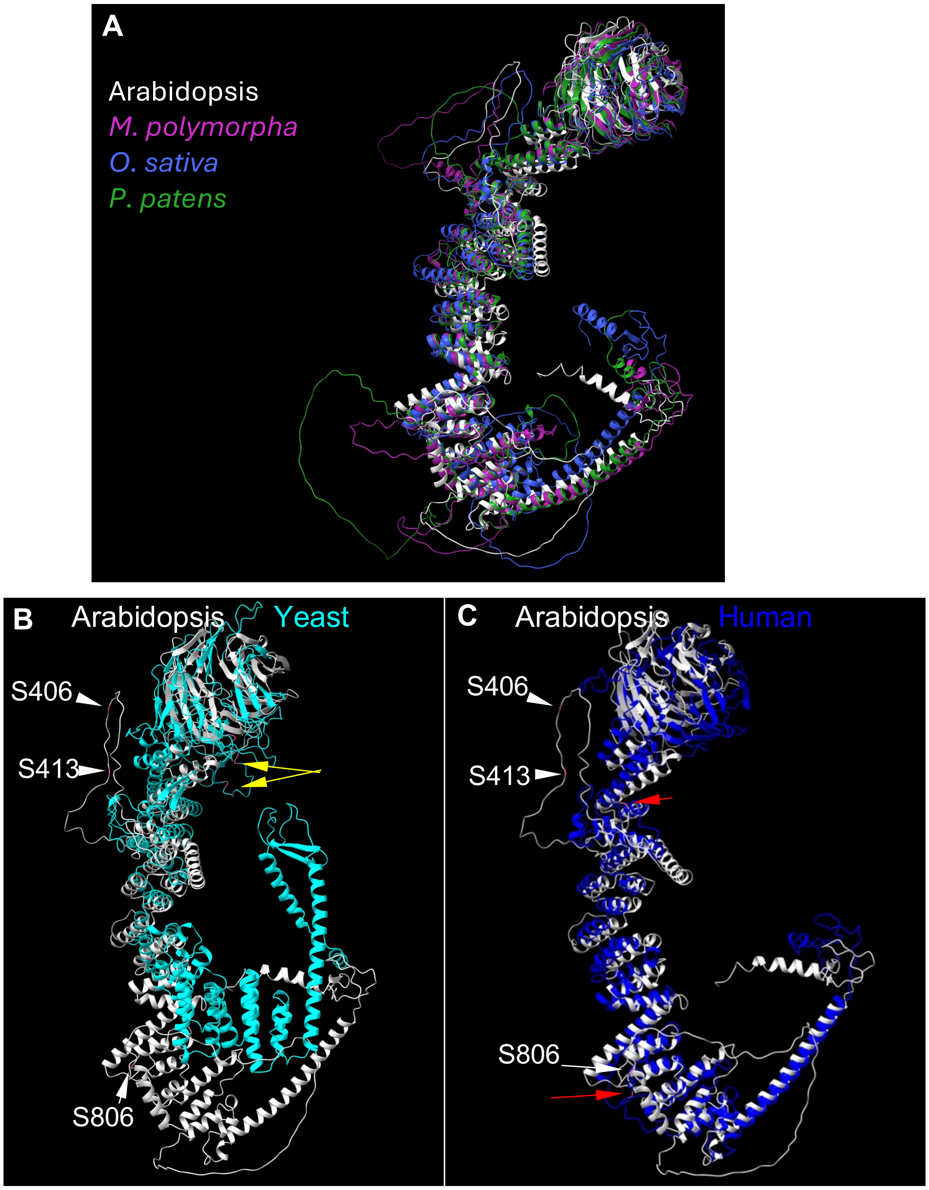


**Supplemental Figure 2. Chimera X protein folding alignments of VPS39.** (A) AlphaFold predictions and alignment of VPS39 homologs from *Marchantia polymorpha* (magenta), *Physcomitrium patens* (green), *Oryza sativa* (blue) and *Arabidopsis thaliana* (gray). All plant VPS39 structures are predicted to have two large unstructured loops similar to Arabidopsis. (B-C) Neither *Sacharomyces cerevisiae* (B) or human (C) VPS39 proteins show similar unstructured domains as Arabidopsis. (B) Arrowheads show position of S406, S413, S806 in Arabidopsis and yellow arrows point to phosphorylated residues in the yeast protein (S246, S247, S249, and S250). C) Red arrow points to S441 and T646 in human VPS39, which have been detected as phosphorylated.

**Supplemental Dataset 1. Details on generalized linear mixed models for stomata assays**

#### Confidence intervals for bootstrap analysis from Generalized linear mixed models for stomata assay, WT vs *vps39-2* (Fig 1F)

Model containing genotype was favored with a p-value of 0.01.

| Biological replicates (n) | # Guard cells | #plants |
|---|---|---|
| WT | 97 | 5 |
| *vps39-2* | 139 | 8 |

| Comparison | 95% confidence interval for bootstrap analysis |
|---|---|
| WT - *vps39-2* | [0.31, 9.66] |

#### Confidence intervals for bootstrap analysis from Generalized linear mixed models for stomata assay, WT vs *vps39-2* vs, *vps39-2* VPS39$^{SD}$-GFP vs *vps39-2* VPS39$^{SA}$-GFP (Fig 3E)

| Biological replicates (n) | # Guard cells | #plants |
|---|---|---|
| WT | 408 | 23 |
| *vps39-2* | 436 | 26 |
| *vps39-2* VPS39$^{SA}$-GFP | 107 | 9 |
| *vps39-2* VPS39$^{SD}$-GFP | 97 | 9 |
| *vps39-2* VPS39-RFP | 214 | 9 |

Model containing genotype was favored with a p-value of $1.304 \times 10^{-7}$. Data from the first stomata experiment was added for the WT and *vps39-2* genotypes to improve the robustness of the model.

| Comparison | 95% confidence interval for bootstrap analysis |
|---|---|
| *vps39-2*-WT(*) | [-3.4, -0.93 ]; |
| *vps39-2* VPS39$^{SA}$-GFP – WT (*) | [-2.12, -0.11 ] |
| *vps39-2* VPS39$^{SD}$-GFP – WT (*) | [-2.13, -0.09 ] |
| *vps39-2* - *vps39-2* VPS39$^{SA}$-GFP (NS) | [-2.27, 0.25 ] |
| *vps39-2* - *vps39-2* VPS39$^{SD}$-GFP (NS) | [-2.31, 0.14 ] |
| *vps39-2* VPS39$^{SA}$-GFP - *vps39-2* VPS39$^{SD}$-GFP (NS) | [-1.18, 1.12 ] |
| *vps39-2* VPS39-RFP – WT (NS) | [-1.1, 0.22] |
| *vps39-2* - *vps39-2* VPS39-RFP (*) | [-2.97, -0.58] |

**Table S1. Comparison of medians of aborted embryos from Figure 3B.** Kruskal-Wallis followed by post-hoc Dunn's test with Šidák correction for multiple comparisons. $p< 0.005116$.

| **Genotype Comparison** | **p-value** |
|---|---|
| WT - *vps39-3* VPS39-GFP | 0.23 |
| WT - *vps39-1/+* | $4.82x10^{-13}$ |
| WT - *vps39-1/+* $VPS39^{SA}$-GFP/+: | $2.52x10^{-6}$ |
| WT - *vps39-1/+* $VPS39^{SD}$-GFP/+: | $8.97x10^{-5}$ |
| *vps39-3* VPS39-GFP - *vps39-1/+*: | $3.32x10^{-8}$ |
| *vps39-3* VPS39-GFP - *vps39-1/+* $VPS39^{SA}$-GFP/+: | $1.50x10^{-3}$ |
| vps39-3 VPS39-GFP- *vps39-1/+* $VPS39^{SD}$-GFP/+: | $1.10x10^{-2}$ |
| *vps39-1/+* - *vps39-1/+* $VPS39^{SA}$-GFP/+: | $1.26x10^{-2}$ |
| vps39-1/+ - *vps39-1/+* $VPS39^{SD}$-GFP/+: | $4.74x10^{-3}$ |
| *vps39-1/+* $VPS39^{SA}$-GFP/+ - *vps39-1/+* $VPS39^{SD}$-GFP/+: | 0.63 |

**Table S2. Pairwise comparison of medians for unfertilized ovules from Figure 3C.** Kruskal-Wallis followed by post-hoc Dunn's test with Šidák correction for multiple comparisons. $p< 0.005116$.

| **Genotype Comparison** | **p-value** |
|---|---|
| WT - *vps39-3* VPS39-GFP | $8.199x10^{-2}$ |
| WT - *vps39-1/+* | $1.59x10^{-2}$ |
| WT - *vps39-1/+* $VPS39^{SA}$-GFP/+: | $2.296x10^{-11}$ |
| WT - *vps39-1/+* $VPS39^{SD}$-GFP/+: | $7.80x10^{-4}$ |
| *vps39-3* VPS39-GFP - *vps39-1/+*: | 0.59 |
| *vps39-3* VPS39-GFP - *vps39-1/+* $VPS39^{SA}$-GFP/+: | $7.352x10^{-6}$ |
| vps39-3 VPS39-GFP- *vps39-1/+* $VPS39^{SD}$-GFP/+: | 0.13 |
| *vps39-1/+* - *vps39-1/+* $VPS39^{SA}$-GFP/+: | $3.67x10^{-5}$ |
| vps39-1/+ - *vps39-1/+* $VPS39^{SD}$-GFP/+: | 0.29 |
| *vps39-1/+* $VPS39^{SA}$-GFP/+ - *vps39-1/+* $VPS39^{SD}$-GFP/+: | $4.51x10^{-3}$ |

**Table S3. Reference phosphopeptides for PHOT1 and OST1.** Guard cell-enriched phosphoproteomics study shows expected phosphorylation in blue light and ABA signaling proteins. Positive Log2FC denotes increased levels of phsphorylation in closed samples. Significant Log2FC > 0.5 or <-0.5. p< 0.05.

| Gene symbol | Araport ID | Residue | Log2FC Closed-Open | p-value Closed-Open |
|---|---|---|---|---|
| PHOT1 | AT3G45780 | S185 | -0.61 | $4.43 \times 10^{-5}$ |
| PHOT1 | AT3G45780 | S350 | -0.94 | 0.01 |
| PHOT1 | AT3G45780 | S410 | -1.86 | $6.01 \times 10^{-18}$ |
| PHOT1 | AT3G45780 | S410 | -1.18 | $5.59 \times 10^{-7}$ |
| OST1 | AT4G33950 | S175; T176 | 2.21 | $4.43 \times 10^{-5}$ |

**Table S4. HOPS, CORVET and vacuolar SNARE proteins detected in proteome.** Proteins are differentially expressed when Log2FC > 0.5 or <-0.5 and p<0.05.

| Gene symbols | Araport ID | AA | Log2FC Closed-Open | p-value Closed-Open |
|---|---|---|---|---|
| VPS3 | AT1G22860 | 984 | 0.18 | 0.15 |
| VPS39 | AT4G36630 | 1000 | 0.07 | 0.21 |
| VPS41 | AT1G08190 | 980 | 0.27 | $4.79 \times 10^{-4}$ |
| VPS11 | AT2G05170 | 932 | 0.27 | $3.74 \times 10^{-4}$ |
| VPS18 | AT1G12470 | 988 | 0.29 | $2.59 \times 10^{-4}$ |
| VPS33 | AT3G54860 | 592 | 0.10 | 0.23 |
| VTI11 | AT5G39510 | 221 | -0.07 | 0.62 |
| SYP22 | AT5G46860 | 268 | -0.14 | 0.03 |
| SYP51 | AT1G16240 | 232 | -0.06 | 0.32 |
| VAMP713 | AT5G11150 | 221 | -0.12 | 0.13 |
| VAMP727 | AT3G54300 | 240 | 0.26 | $2.83 \times 10^{-3}$ |

**Table S5. Detected phosphorylation of HOPS, CORVET and vacuolar SNARE proteins.** Significantly enriched phosphopeptides are those with Log2FC > 0.5 or <-0.5 and p-value <0.05.

| Gene symbols | Araport ID | PTM protein localization | Missed cleavages | Log2FC Closed-Open | p-value Closed-Open |
|---|---|---|---|---|---|
| VPS39 | AT4G36630 | S392 | 0 | -0.09 | 0.56 |
| VPS39 | AT4G36630 | S413 | 0 | 0.79 | $5.41x10^{-6}$ |
| VPS39 | AT4G36630 | S806 | 0 | 0.40 | $2.102x10^{-3}$ |
| VPS39 | AT4G36630 | S806 | 1 | -0.08 | 0.38 |
| VPS41 | AT1G08190 | S859/S860 | 2 | 0.2 | 0.18 |
| VPS41 | AT1G08190 | S860 | 1 | -0.02 | 0.88 |
| VPS41 | AT1G08190 | S860 | 2 | 0.52 | 0.08 |
| VTI11 | AT5G39510 | S25 | 1 | -0.1 | 0.56 |
| VTI11 | AT5G39510 | S79 | 2 | -0.31 | 0.06 |
| VAMP727 | AT3G54300 | S194 | 0 | -0.24 | 0.07 |

**Table S6. Primers used in this study.**

| **Primer Name** | **Sequence 5'-3'** | **Purpose** |
|---|---|---|
| VPS39-4R | GCTACCGTCTTCTCTAGCACGT | *vps39-2* genotyping |
| Wisc-DS-Lox-p745 | AACGTCCGCAATGTGTTATTAAGTTGTC | *vps39-2* genotyping |
| VPS39UTR-F | GGTTATTCTCATTCATAGAAGAAAAGCCA | *vps39-2* genotyping |
| PEX4-q1F | CTT AAC TGC GAC TCA GGG AAT CTT CTA AG | qPCR Housekeeping gene |
| PEX4-q1R | TCA TCC TTT CTT AGG CAT AGC GGC | qPCR Housekeeping gene |
| PUX7-qF1 | CTCCTCATTCCAAACCCAAAGAGGA | qPCR Housekeeping gene |
| PUX7-qR1 | AAGCAGCTAATGCTCGTTGCAG | qPCR Housekeeping gene |
| VPS39-Q22F | GGTGGGAGAGGATCAACGGT | qPCR Target gene |
| VPS39-Q23R | AAACGGAAGCAGGTTATGGAGT | qPCR Target gene |
| LBb1.3 | ATTTTGCCGATTTCGGAAC | *vps39-1* genotyping |
| VPS39-5R | CACTCTGCCTCAAACTCTTGATGACTG | *vps39-1* genotyping |
| vps39-1 F1 | CTTTGTTGAAGCGTCTTCCAC | *vps39-1* genotyping and phosphomutant genotyping and sequencing |
| VPS39 Term-R10 | CAGACAAACAACACGAAAGGAG | *vps39-1* genotyping and phosphomutant genotyping and sequencing |
| GFP R1 | GTCGTGCTGCTTCATGTGG | phosphomutant genotyping and sequencing |
| VPS39-RP-GABI | TTGAACTCTCTTTCGCAGCTC | *vps39-3* genotyping |
| GABI-LB | ATATTGACCATCATACTCATTGC | *vps39-3* genotyping |
